# The Role of Scale in the Estimation of Cell-type Proportions

**DOI:** 10.1101/857805

**Authors:** Gregory J. Hunt, Johann A. Gagnon-Bartsch

**Affiliations:** DEPARTMENT OF MATHEMATICS, WILLIAM & MARY; DEPARTMENT OF STATISTICS, UNIVERSITY OF MICHIGAN

## Abstract

Complex tissues are composed of a large number of different types of cells, each involved in a multitude of biological processes. Consequently, an important component to understanding such processes is understanding the cell-type composition of the tissues. Estimating cell type composition using high-throughput gene expression data is known as cell-type deconvolution. In this paper, we first summarize the extensive deconvolution literature by identifying a common regression-like approach to deconvolution. We call this approach the Unified Deconvolution-as-Regression (UDAR) framework. While methods that fall under this framework all use a similar model, they fit using data on different scales. Two popular scales for gene expression data are logarithmic and linear. Unfortunately, each of these scales has problems in the UDAR framework. Using log-scale gene expressions proposes a biologically implausible model and using linear-scale gene expressions will lead to statistically inefficient estimators. To overcome these problems, we propose a new approach for cell-type deconvolution that works on a hybrid of the two scales. This new approach is biologically plausible and improves statistical efficiency. We compare the hybrid approach to other methods on simulations as well as a collection of eleven real benchmark datasets. Here, we find the hybrid approach to be accurate and robust.deconvolution, gene expression, microarray, RNA-seq

## 1. INTRODUCTION

The tissues of multi-cellular organisms are typically comprised of a combination of many types of cells. As each cell type has its own set of functions and behaviors, the composition and interaction of different cell types is integral to the function and behavior of the tissues. Thus, studying cell-type composition has long been of broad biological interest. Examples of the importance of cell type composition abound from the biological literature. In the study of infectious diseases, the composition of white blood cells is important as it is indicative of many types of dysfunctions (George and Panos, 2007). For example, the number of T-cells among human peripheral blood mononuclear cells (PBMCs) spikes after a Lyme infection (Bouquet *et al.*, 2016). In neuroscience, the composition of brain cells has long been a subject of study. For example, studying the relative composition of microglia in human brains is of interest for those studying developmental dynamics (Ayana *et al.*, 2018). Similarly, understanding changes in the number of neuron and glial cells has been the subject of extensive study with regards to Alzheimer’s disease (Mohammadi *et al.*, 2015).

For this reason, methods to estimate cell-type proportions from high-throughput genomics data have been extensively studied over the past two decades (for comprehensive literature reviews see Gaujoux (2013) or Mohammadi *et al.* (2015)). Estimating cell-type proportions is known as *cell-type deconvolution*. Given gene expression data from sample comprised of a mixture of cell types, deconvolution methods estimate the proportions of the constituent cell types. These cell-type proportions may be of interest in their own right, for example, to track the changes in cell-type composition over time (Newman *et al.*, 2015). In other cases, the estimated cell-type proportions are used as a means of deconfounding differential expression analysis (Capurro *et al.*, 2015). In this case, the cell-type proportions can help explain observed gene-expression differences across samples. By including the estimated proportions in a model, one can separate differences coming from within-cell-type changes in gene expression and those differences coming purely from cell-type-compositional differences among samples (Hagenauer *et al.*, 2016).

In this paper, we present a critique of existing cell-type deconvolution methods and present a new method for cell-type deconvolution that addresses the issues we raise. First, in Section 2, we characterize existing deconvolution literature, proposing a new unified deconvolution framework called the Unified Deconvolution-as-Regression (UDAR) framework. The UDAR framework summarizes much of the existing deconvolution literature, including many popular deconvolution methods. It demonstrates that these methods employ a common unified model of the data and mainly differ in how their parameter estimates are fit. One important fitting consideration is data scale. Broadly, methods either fit using linear-transformed or log-transformed gene expression data. Unfortunately, each of these scales has problems. We will show that using log-scale gene expressions proposes a biologically implausible model and that using linear-scale gene expressions will lead to statistically inefficient estimators. Using the UDAR framework as a point of comparison, in Section 3 we introduce a hybrid-scale approach for cell-type deconvolution that uses aspects of fitting on both scales. Subsequently, in Section 4, we evaluate the performance of the new approach across a wide range of simulated data as well as eleven real benchmark datasets. These comprehensive analyses show that the proposed approach produces accurate and robust estimates of cell-type proportions.

## 2. A UNIFIED FRAMEWORK FOR EXISTING DECONVOLUTION MODELS

Let *Y* ∈ ℝ^*N*^ be the measurements of *N* gene expressions in a mixture sample of *K* types of cells and *R* ∈ ℝ^*N*×*K*^ be reference expressions of these *N* genes across the *K* constituent cell types. Furthermore, let *p* = (*p*_1_, …, *p*_*K*_) be the proportions of the *K* cell types in the mixture sample. Implicit in being proportions is that *p* must satisfy the sum-to-one (STO) constraint: 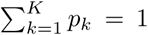, and the non-negativity (NN) constraint: *p*_*k*_ ≥ 0 for *k* = 1, …, *K*. That is, *p* ∈ Δ_*K*−1_, the (*K* − 1) probability simplex Δ_*K*−1_ = {*x* ∈ ℝ^*K*^ : *x*_*k*_ ≥ 0 and 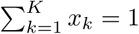}.

The deconvolution problem is that *p* is unknown and we want to estimate it. In this section we introduce a new unified model for cell-type deconvolution called the Unified Deconvolution-as-Regression (UDAR) framework. The UDAR framework posits that *Y, R* and *p* are related through the linear model

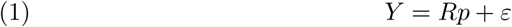

for a random error *ε*. Estimating *p* under this model is equivalent to solving a constrained regression of *Y* on *R* where the coefficients *p* must live in Δ_*K*−1_. Hence, using this model is treating deconvolution as regression. Note that we only consider problems where *Y* and *R* are known and we are interested in estimating *p*. We do not consider the related problem where *R* is also unknown. For a discussion of this problem see Gaujoux (2013) or Mohammadi *et al.* (2015) or Wang *et al.* (2016).

Nearly all existing methods model deconvolution following Equation (1), however a few exceptions exist, e.g. Hunt *et al.* (2019). What differs among the methods is the approach by which *p* is estimated. There are common themes among how estimates of 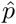 are fit. Typically, methods specify: (1) a loss function *L* : ℝ^*K*^ → ℝ_+_ that determines model fit *L*(*p*) for putative proportions *p*, (2) an optimization space Π ⊆ ℝ^*K*^ for *p*, and (3) a post-hoc adjustment function *φ* : Π → Δ_*K*−1_ mapping from the optimization space Π to the desired simplex Δ_*K*−1_. They then estimate *p* by minimizing *L* over Π and applying *φ*. This approach is described in in Algorithm 1.

### Algorithm 1 UDAR Fitting

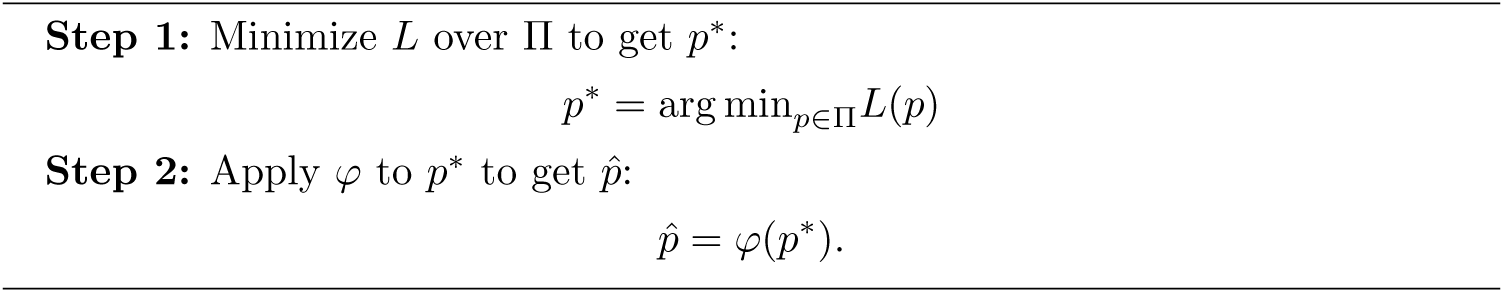

The idea behind this approach is that, while ideally 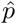 is the minimizer of *L* over Δ_*K*−1_, solving such a constrained minimization problem is difficult. Thus, UDAR methods solve an easier relaxation of this problem, minimizing *L* over Π ⊇ Δ_*K*−1_ and then making post-hoc adjustments to *p*\* to produce a final estimate 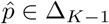.

A large number of existing deconvolution methods fit into this framework under appropriate choices of *L*, Π, and *φ*. The most common choice of loss is the squared-error loss (Lu *et al.*, 2003; Abbas *et al.*, 2009; Wang *et al.*, 2006; Gong *et al.*, 2011; Qiao *et al.*, 2012; Racle *et al.*, 2017). Other loss functions used include an elastic net penalized loss (Altboum *et al.*, 2014), a support-vector regression approach which is equivalent to using an *ϵ*-insensitive loss (Newman *et al.*, 2015), and a Bayesian-likelihood approach based on Latent Dirichlet Allocation that is equivalent to letting *L* be a likelihood-based loss (Qiao *et al.*, 2012; Blei *et al.*, 2003). The optimization space Π is typically one of three spaces: (1) Δ_*K*−1_ (Gong *et al.*, 2011), (2) 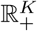, the positive orthant of ℝ^*K*^ (Qiao *et al.*, 2012; Racle *et al.*, 2017), or (3) ℝ^*K*^ (Lu *et al.*, 2003; Abbas *et al.*, 2009; Wang *et al.*, 2006; Newman *et al.*, 2015). In the first case where Π = Δ_*K*−1_ no post-hoc adjustments are necessary and so *φ* is the identity function. In the second case where 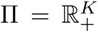, since *p*\* already satisfies the NN constraint *φ* re-normalizes *p*\* to enforce the STO constraint and hence 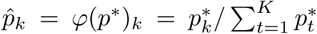 (Qiao *et al.*, 2012; Racle *et al.*, 2017). Finally, in the first case of unconstrained optimization where Π = ℝ^*K*^, *φ* zeros out negative coefficients and then re-normalizes so that 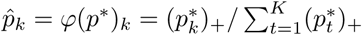 where (·)_+_ = max(., 0) is the positive part (Lu *et al.*, 2003; Abbas *et al.*, 2009; Wang *et al.*, 2006; Newman *et al.*, 2015). We call this latter post-hoc adjustment the “zero-then-renormalize” adjustment.

### 2.1. Scale Considerations for Deconvolution

An important question for the UDAR framework is the appropriateness of this model for the deconvolution problem. One important modeling consideration is data scale. Typically, gene expressions are either linearly transformed, e.g. TPM (Conesa *et al.*, 2016), or logarithmically transformed, e.g. RMA, (Irizarry *et al.*, 2003). In the former case, we say the data is on the linear-scale and in the latter we say the data is on the log-scale. Some deconvolution methods assume linear-scale expressions like in Newman *et al.* (2015), some methods assume log-scale expressions as in Qiao *et al.* (2012), most make no explicit assumptions about data scale at all. In the following sections we will consider the appropriate data scale for the UDAR model. This will primarily concern the two major components of the model: (1) the linear mean-structure *Rp* and (2) the additive error-structure *ε*.

#### 2.1.1. Mean Modeling

Assume we have a mixture sample comprised of cell types *k* = 1, …, *K* in proportions *p*_1_, …, *p*_*K*_. First, notice that if *η*_*n*_ is the amount of mRNA in our mixture sample coming from gene *n* and *η*_*nk*_ is the amount of that mRNA in the sample coming from type *k* cells then,

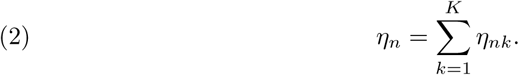

Now, assume we also have some reference sample of type *k* cells. Let the amount of mRNA from gene *n* in the reference sample be 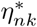. Since the mixture sample is comprised of a proportion *p*_*k*_ of type *k* cells and the reference sample is 100% type *k* cells, then we expect that

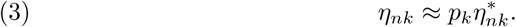

Essentially, this assumes that type *k* cells in the mixture behave as if they were a random sample of the type *k* reference cells. We assume that this relationship is only approximate because the type *k* reference cells may not exactly mimic the type *k* mixture cells. For example, the microenvironment of the cells in the mixture may modify gene expression.

Combining Equations (2) and (3), we get that

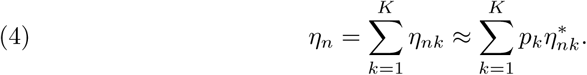

Now assume that the *linear-scale* measured gene expressions are proportional to the amount of mRNA so that *Y*_*n*_ ≈ *γα*_*n*_*η*_*n*_ and 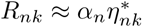 for constants *γ* and {*α*_*n*_}. The proportionality constants *α*_*n*_ capture gene-specific effects like probe affinity (for microarray data) or length-biases (for RNA-seq). The multiplier *γ* captures global differences between the mixture and references. This includes effects like sequencing depth or amount of mRNA. Again, we assume approximate equality because the measurement process may introduce random errors. Combining with Equation (4) we now get that

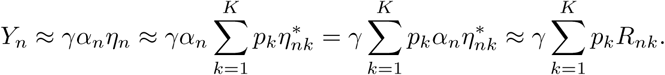

As is customary, assume that *Y* and *R* have been normalized to account for global expression differences, by e.g. TPM (Conesa *et al.*, 2016), so that *γ* = 1. Then the above equation shows that the linear model *Y* ≈ *Rp* proposed by UDAR is correctly specified for linear-scale gene expression measurements since 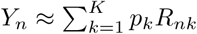.

However, even if *γ* = 1, the linear mean structure is mis-specified for log-scale gene expressions as

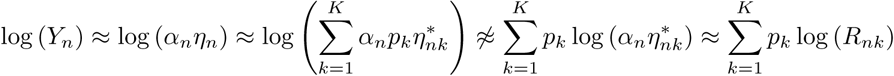

since we can’t interchange a sum and a log. Thus log 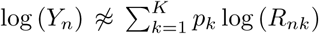 and so a linear mean-structure as proposed by UDAR does not make sense on the log-scale. For a toy example of this principle see Figure 1.

**FIGURE 1.**
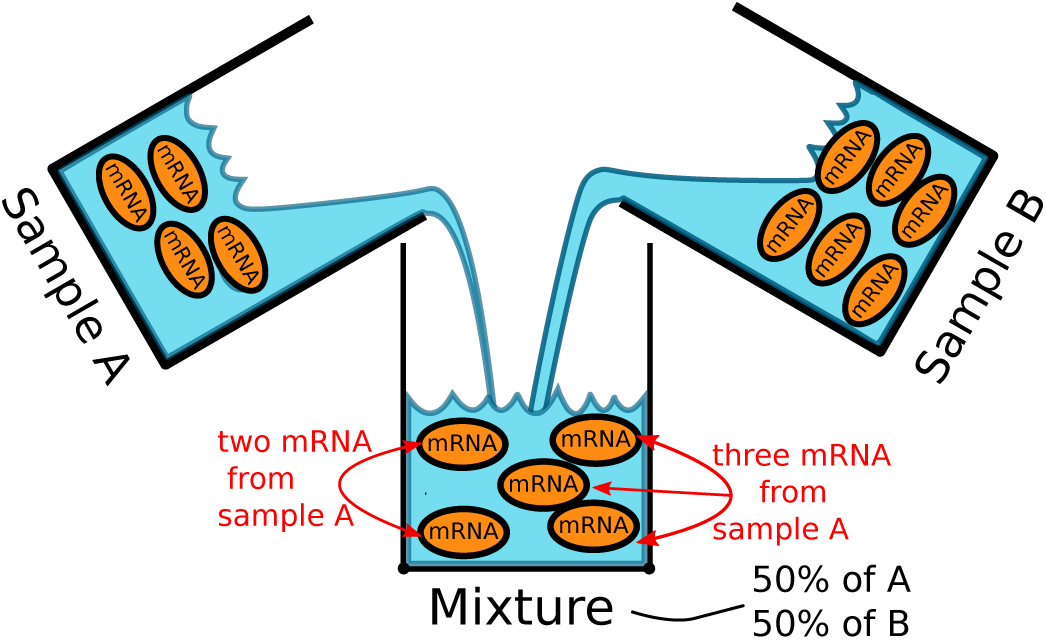
The mixture sample is 50% of A and 50% of B. The orange ovals represent mRNA from a specific gene. Since the reference of type A typically has four mRNA we expect 4 ×.5 = 2 mRNA in the mixture to come from ref. A. Similarly, since ref. B typically has six mRNA, we expect 6 × .5 = 3 mRNA in the mixture to have come from ref. B. In total, we get 5 = 4 × .5 + 6 × .5 mRNA in the mixture. Thus the amount of mRNA in the mixture is a linear mixture of the amount of mRNA. This does not work if we logarithmically transform the counts. In that case we would expect, on the log-scale, to get *log*(4) × .5 + log(6) × .5 ≈ 1.6 mRNA. Exponentiating back to the linear scale, this is ≈ 4.9, thus under-counting the true amount of mRNA.

#### 2.1.2. Error Modeling

In contrast to the mean-structure, error assumptions are most reasonable for log-scale expressions. While most methods simply note that *Y* ≈ *Rp* and do not explicitly include an error term *ε* in their models, their loss functions are optimal for typical regression-like error assumptions about *ε*. For example, deconvolution methods minimizing the squared-error loss are are optimal when 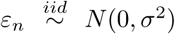 with some constant error variance *σ*^2^ > 0. Such regression assumptions are most appropriate on the log-scale. Indeed, it has been widely noted that errors are well-modeled as normal with approximately constant variance across expression levels for log-scale gene expression data (Qiao *et al.*, 2012). Conversely, error for linear-scale expression data are right-skewed and the variance tends to increase with increasing mean expression (Qiao *et al.*, 2012; Hardin and Wilson, 2009; Weng *et al.*, 2006; Tu *et al.*, 2002; Zwiener *et al.*, 2014).

## 3. A HYBRID MODEL FOR DECONVOLUTION

The previous two sections present a problem for existing deconvolution methods. If they follow the UDAR model on the log-scale, they will have a mis-specified mean. Conversely, if they propose the UDAR model with linear-scale expressions, the error assumptions are un-realistic. To avoid both of these problems, we propose a new method based on a hybrid of the two scales. Our hybrid model proposes that

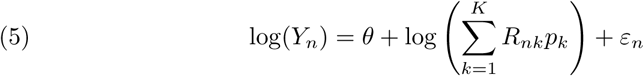

where 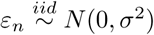. Again, *Y*_*n*_ and *R*_*nk*_ are the linear-scale gene expression. Our hybrid-scale model proposes additive Gaussian error *after* a log transformation and thus uses an appropriate scale for errors. Furthermore, the mean-structure in Equation 5 implies log(*Y*) ≈ *θ* + log(*Rp*), or equivalently, *Y* ≈ *e*^*θ*^*Rp*. Thus it proposes a plausible linear-mixing structure on the *linear-scale* as discussed in Section 2.1.1. Indeed, *e*^*θ*^ is precisely the term *γ* mentioned in this section. Thus the hybrid model explicitly includes a term that allows it to naturally account for systematic differences between the mixture and reference expressions.

### 3.1. Fitting The Hybrid Model

To estimate *p* under this model, we let 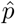 be the MLE so that

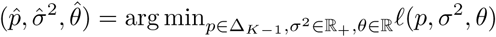

and *ℓ* is the joint log-likelihood function of *p, σ*^2^, and *θ*. A valuable property of this hybrid model is that this optimization problem can be easily solved using an approach analogous to the UDAR fitting procedure.

Define 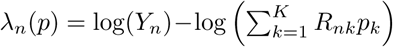 and let *S*^2^(*p*) be the sample variance of the *λ*_*n*_(*p*),

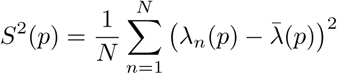

where 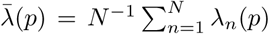. It can be shown (see Supplementary Section 1) that 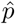 is the minimizer of *S*^2^ so that

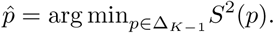

Furthermore, since *S*^2^ is invariant under scaling so that *S*^2^(*cp*) = *S*^2^(*p*) for any *c* ∈ ℝ_+_ we do not need to optimize over Δ_*K*−1_ directly. Instead, we can solve this optimization problem over any set containing Δ_*K*−1_ and simply re-normalize. Let *p*\* be any minimum of *S*^2^(*p*) over *p* ∈ Π where Π is any set satisfying 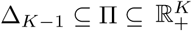 so that

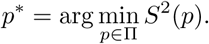

(Notice that this minimum is not unique since if *p*\* minimizes *S*^2^ then so does *cp*\*.) Then if 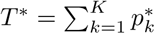 is the sum of the elements of *p*\*, the MLE for *p* is 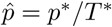. Full details for this fact can be found in Supplementary Section 1. This motivates Algorithm 2, a simple procedure to estimate 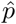.

#### Algorithm 2 Hybrid Fitting Procedure

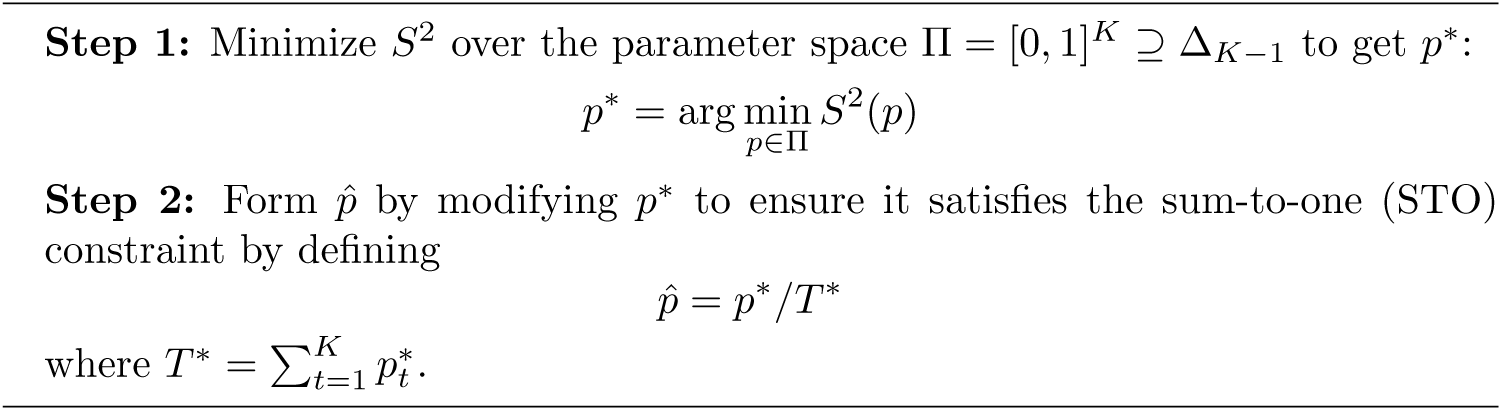

This procedure allows us to find the MLE without trying to minimize *ℓ* over *p, σ*^2^ and *θ* simultaneously. Furthermore, like the UDAR model, this procedure also allows us to optimize *p* over a relaxation Π = [0, 1]^*K*^ ⊇ Δ_*K*−1_ instead of having to directly search over Δ_*K*−1_. Also, like Algorithm 1, we re-normalize *p*\* to form a 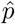 that is in Δ_*K*−1_. However, while for the UDAR framework the post-hoc adjustments were a heuristic to enforce constraints on 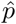, our re-normalization is not heuristic. The two steps in Algorithm 2 precisely allow us to recover the MLE of the hybrid model without solving a difficult optimization problem over Δ_*K*−1_. While we could have optimized *S*^2^(*p*) over any space Π where 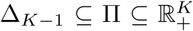, letting Π = [0, 1]^*K*^ greatly simplifies the optimization problem and allows us to use standard global optimization routines with box constraints to find 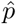.

In this section we considered how to estimate *p* as the MLE. Here, we showed that finding the MLE reduces to minimizing a variance-based loss *S*^2^(*p*). More generally, one could use the model in Equation 5 and estimate *p* by minimizing other loss functions. For example, one could consider L1 or L2 penalized losses, or an *ε*-insensitive loss. Thus, the model proposed by Equation 5 is general and extensible in many of the same ways as the UDAR framework.

### 3.2. References, Marker Genes, and Weights

We are considering the deconvolution problem where there is some known reference data available. This data is typically obtained from online gene expression repositories like GEO (Edgar, 2002) or from specific profiles complied for cell-type deconvolution.

Such reference data is used in two major ways. First, the reference data is used to create the reference matrix *R* so that *R*_*nk*_ is the typical expression of gene *n* in a sample purely of cells of type *k*. Often, there exists more than one reference sample for a particular cell type. If one has *ν*_*k*_ reference profiles of cell type *k*, then *R*_*nk*_ is typically average expression across the profiles so that 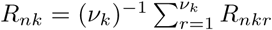 where *R*_*nkr*_ is the gene expression of gene *n* in the *r*^*th*^ reference of cell type *k*.

In addition to using reference data to form the reference matrix *R*, this reference data is often used to find marker genes. Marker genes are genes that are particularly highly expressed in one cell type but not the others. Typically marker genes are identified by comparing gene expression across cell types in the reference data using, for example, a *t*-test. Once identified, deconvolution methods fit using only the subset of marker genes. Let ℳ ⊆ {1, …, *N*} be the set of marker genes. Then the use of marker genes can be viewed as variable selection where we only fit using those *n* ∈ ℳ. Alternatively, we can view the marker genes as a weighting of the loss function. Under the UDAR model, fitting using ℳ is equivalent to using a weighted loss function with weights *w*_*n*_ = 𝟙(*n* ∈ ℳ).

Our fitting approach in Algorithm 2 can also encompass marker genes as variable selection or a weighted loss. For example, we can let 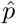 be the MLE of *p* obtained by minimizing *S*^2^(*p*) over only those *n* ∈ ℳ. Equivalently, we can calculate *S*^2^(*p*) as a weighted sample variance with weights *w*_*n*_ ∝ 𝟙(*n* ∈ ℳ). In the case where we have more than one reference of each cell type we may extend this approach by relaxing the equal variance assumption of our model, estimating the variances of the marker genes individually, and weighting inversely accordingly to these variances. This allows us to incorporate variance information from the reference data if it is available.

## 4. RESULTS

### 4.1. Comparison of Methods on Simulated Data

To evaluate the efficacy of the hybrid approach as compared to the UDAR model we evaluate the methods on simulated heterogeneous mixtures of cells. We simulate the mixtures using reference RNA-seq profiles of brain, liver and muscle cells from Parsons *et al.* (2015). First we define *R* ∈ ℝ^*N*×*K*^ as the reference profile matrix of the *N* = 23, 459 genes profiled in the *K* = 3 reference samples (Parsons *et al.*, 2015). Thus *R* is comprised of linear scale (untransformed) read counts. We then generate mixture proportions *p* uniformly from Δ_*K*−1_ and form a simulated mixture profile *Y* ∈ ℝ^*N*^ so that

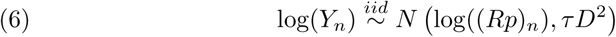

where *D*^2^ is the sample variance of the vectorized log-scale reference matrix and *τ* is a variance multiplier parameter we are free to choose.

To estimate the proportions from this simulated data, we consider four approaches. First we use the hybrid approach outlined in Algorithm 2. We compare this to two simple regression approaches and one more sophisticated support-vector regression approach from the literature, cibersort (Newman *et al.*, 2015). All three of the approaches against which we compare follow the UDAR framework to solve the regression problem. The simple regression approaches follow the UDAR model using a squared-error loss *L*, optimizing over Π = ℝ^*K*^, and applying the simple zero-then-renormalize post-hoc adjustments to ensure the estimates live in Δ_*K*−1_. The simple form of *L* and Π mean that these approaches are equivalent to modified regressions on linear and log-scale data, respectively. We call the linear-scale version a “Regression” approach because it is equivalent to letting 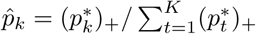 where *p*\* are the coefficients obtained from regressing *Y* on *R*. We call the log-scale version approach “LogRegression” because it is equivalent to the regression approach but where the *p*\* are the coefficients obtained from regressing log(*Y*) on log(*R*). The fourth method, cibersort, also falls under the UDAR framework using an *ε*-insensitive loss (i.e. using support-vector regression) and optimizing over Π = ℝ^*K*^. It then applies the zero-then-renormalize post-hoc adjustment to the estimated parameters. For all four methods we subset *Y* and *R* to a set of *M* ≪ *N* marker genes chosen by an ANOVA on the reference data. The exact same set of marker genes are used to fit the four methods.

In Figure 2 we plot scatter plots of the estimates against the truth for the hybrid approach, the two regression approaches, and cibersort. There are four sub-plots for four different simulation settings. The simulation settings cover a low amount of noise (*τ* = 1*/*4), a large amount of noise (*τ* = 1), and a low number of markers (*M* = 10) and a large number of markers (*M* = 100). For each setting and method we estimate the proportions for 500 simulated samples. From these plots we an see that the hybrid-scale approach generally out-performs the other approaches. The LogRegression approach does comparatively poorly because it has a mis-specified mean and thus an obvious bias manifested in the *S*-shaped relationship between the truth and the estimates. The other three approaches do not exhibit this bias. On average, their estimates generally track the true mixing proportions. Instead of the *S*-shaped curve, we see that for these methods the points in Figure 2 have a scatter centered around the dotted line. Nonetheless, linear-scale regression and cibersort both perform worse than the hybrid approach because they have higher variance estimates. This is evidenced from the higher scatter of the estimates around the dotted line in Figure 2. Thus, the estimates for the hybrid approach are typically closer to the truth than for the other two linear-scale methods (Regression and cibersort). The higher variance of Regression and cibersort follows from the fact that they mis-specify the error scale leading to statistically inefficient estimates. In Supplementary Figure 1 we display boxplots of errors for the four methods over a larger range of simulation settings. These plots show largely the same story observed in Figure 2.

**FIGURE 2.**
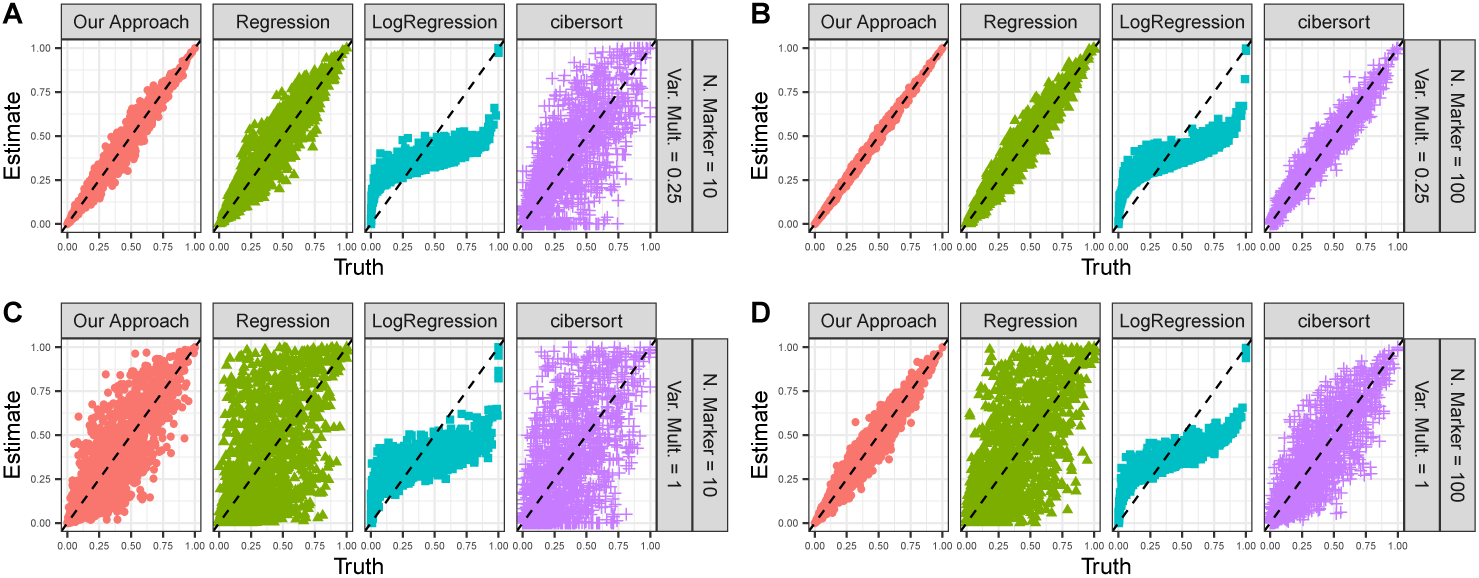
Evaluation of methods on simulated mixture data with Gaussian noise for different values of the variance multiplier (*τ*) and number of marker genes (*M*). (A) *τ* = .25, *M* = 10, (B) *τ* = .25, *M* = 100, (C) *τ* = 1, *M* = 10, (D) *τ* = 1, *M* = 100.

To explore the role of the Gaussianity assumption on performance, in Figure 3 we construct similar scatter plots for data simulated using a negative binomial model for the expressions. The simulations are similar to those in Equation 6, however we let

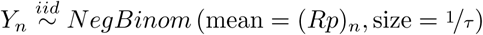

so that *Y*_*n*_ has mean *µ* = 𝔼 [*Y*_*n*_] = (*Rp*)_*n*_ and variance Var (*Y*_*n*_) = *µ* + *µ*^2^*τ*. In Figure 3 we again consider simulation settings for low error (*τ* = ¼), high error (*τ* = 1) and for a small number of markers (*M* = 10) and a larger number of markers (*M* = 100). We see similar behavior for the negative binomial simulations as in the Gaussian case. The hybrid approach out-performs the other approaches, suggesting that the model is relatively insensitive to an exact Gaussian error assumption. The hybrid-scale approach performs better than the other approaches because it uses reasonable scales for both the mean structure and the errors. This leads to both a lower bias and lower variance estimates than the other methods. In Supplementary Figure 2 we display boxplots of errors for the negative binomial simulations over a wider range of simulation settings. These figures tell much the same story.

**FIGURE 3.**
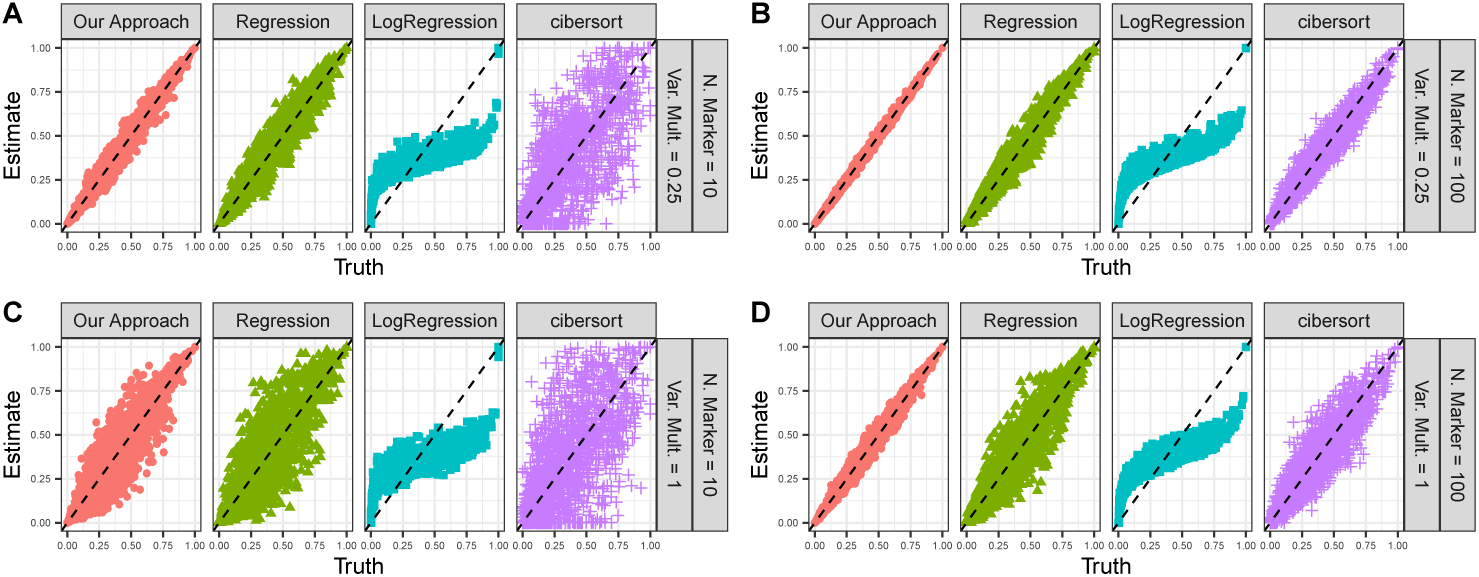
Evaluation of methods on simulated mixture data from a negative binomial model for different values of the variance multiplier (*τ*) and number of marker genes (*M*). (A) *τ* = .25, *M* = 10, (B) *τ* = .25, *M* = 100, (C) *τ* = 1, *M* = 10, (D) *τ* = 1, *M* = 100.

### 4.2. Comparison of Methods on Real Data

To evaluate the performance of the four deconvolution methods on real data we use a collection of existing deconvolution benchmark datasets (see Supplementary Table 1). In all, these eleven datasets cover a range of realistic deconvolution settings. Across the datasets there is a range of cell types, number of cell types, organisms (human and rat), and technologies (RNA-seq and microarrays). Some datasets contain reference data created as part of the same sequencing experiment, while other datasets contain third-party references. For most of the datasets, the true mixing proportions are known because the cells were mixed in known proportions before expressions were assayed. However for three of the datasets the true proportions are the cell type proportions reported by a physical sorting technique applied after the gene expression assays.

The choice of marker genes is an extremely important component in the application of cell type deconvolution methods as accuracy is strongly influenced by the choice of markers. For example, consider Figure 4. In this figure we plot the error for the dataset from Gong *et al.* (2011) for the four methods. Error is measured as absolute value of the difference between the true proportions and their predictions. We use the exact same marker genes for each method, but estimate the cell type proportions using a range of different numbers of markers (*M*). We let *M* vary following an approximate exponential sequence *M* = 1, 2, 5, 10, 20, 50, 100, 200, 500, 1000, 2000, 5000, approximately doubling the number of marker genes at each step. We can see from this figure that estimation performance depends heavily on the number of marker genes used. For example, cibersort does poorly for a small number of markers, but sees improvement for a large number of markers. For each method, there is an optimal number of marker genes to use. For the log-scale regression and cibersort this is about 500, for the hybrid approach this is about 100, and for the linear-scale regression, performance appears to improve as we include more marker genes.

**FIGURE 4.**
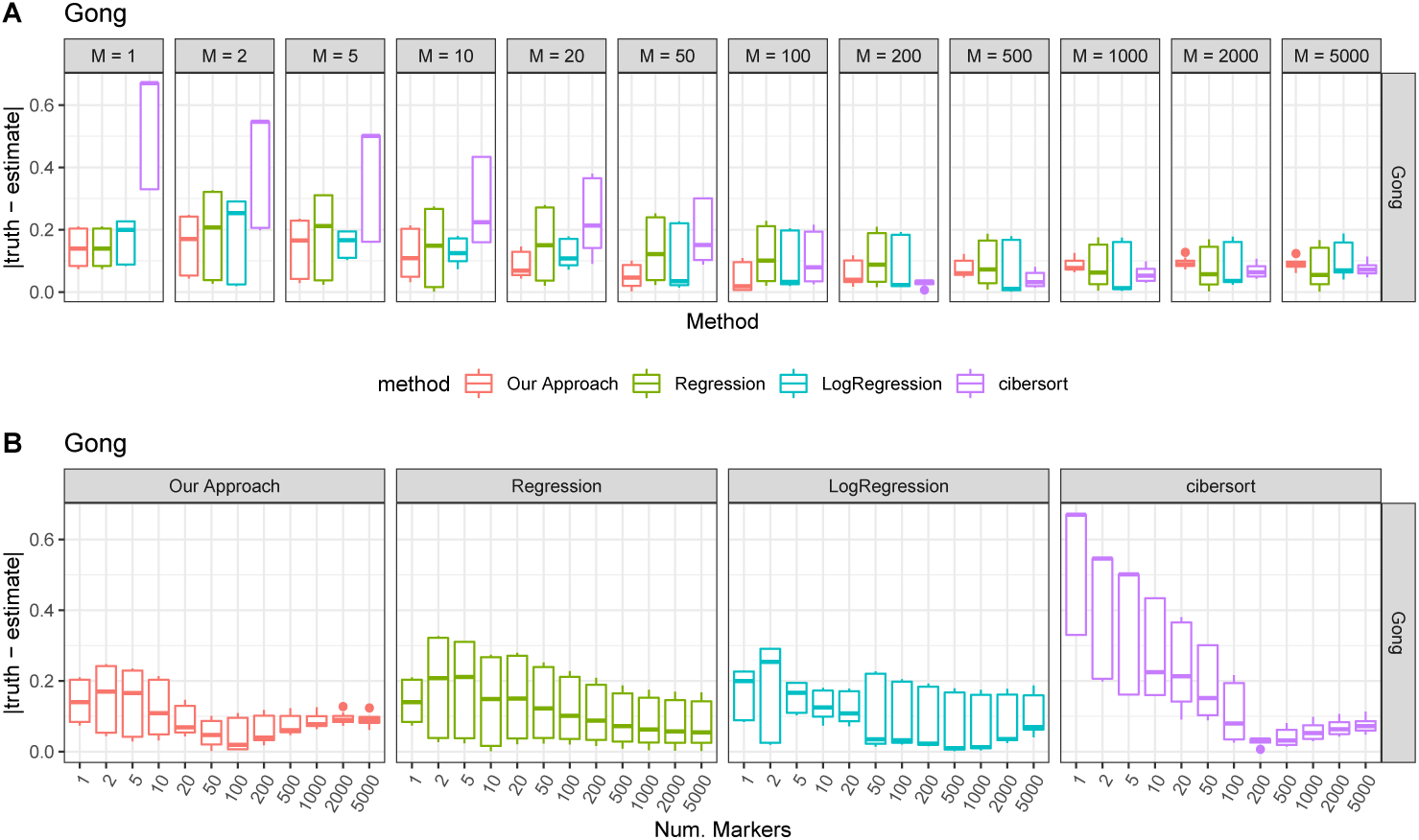
Error for methods for the Gong dataset over a varying number of markers (*M*). Error is measured as the absolute value of the truth less the estimate. (A) displays the plots by number of markers. (B) displays the exact same data but separating by method.

Importantly, the optimal number of marker genes depends as much on the particular dataset as on the particular method. As an example, consider accuracy as a function of number of marker genes in Figure 5 for the data from Shi et al. (MAQC, 2006). Here, error seems to decrease for all methods as we increase the number of marker genes. For this dataset the plot suggests that we should use thousands of marker genes for deconvolution. We display similar plots for the other datasets in Supplementary Figures 3-11. These show that the optimal number of marker genes varies widely from dataset to dataset and method to method.

**FIGURE 5.**
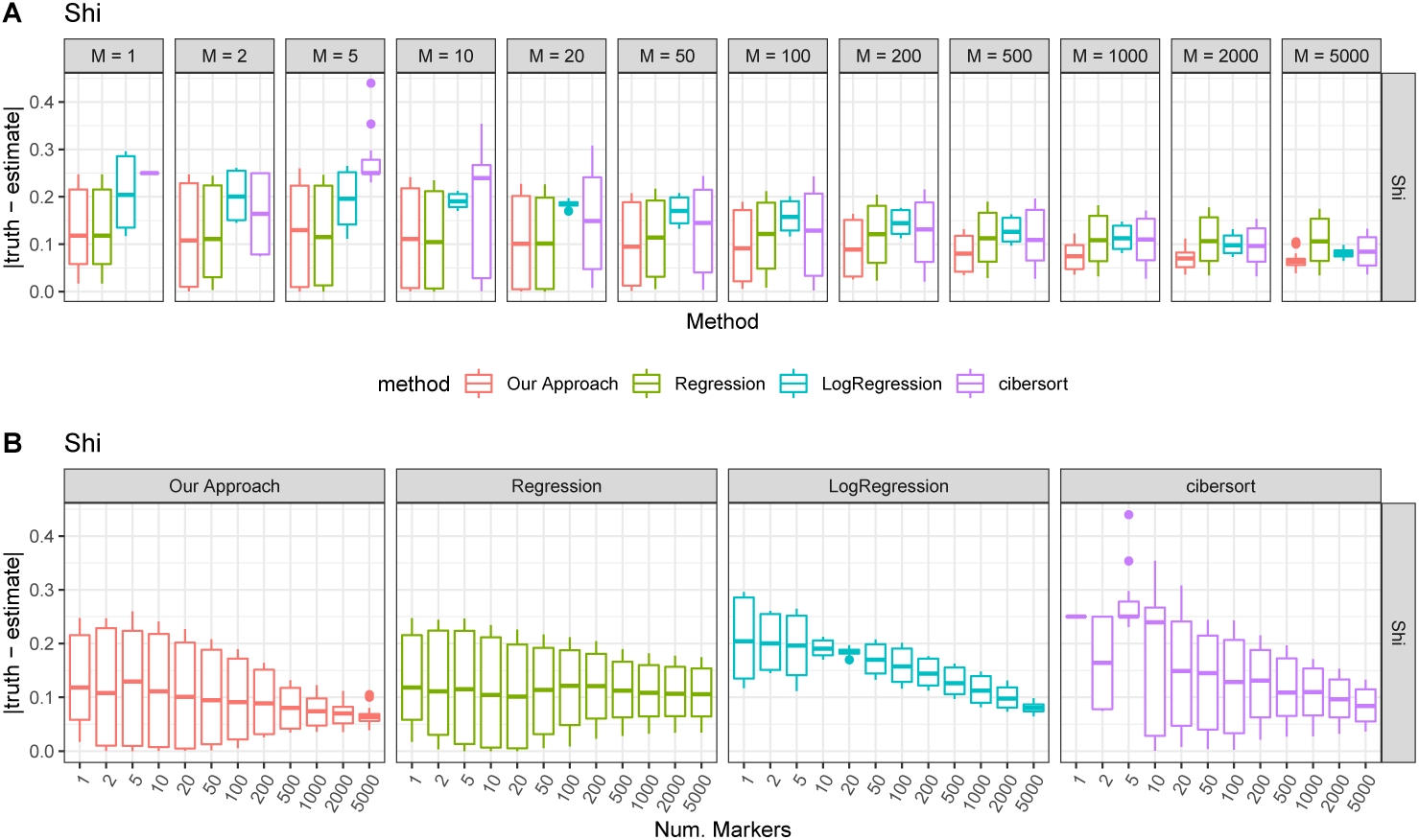
Error for methods for the Shi dataset over a varying number of markers (*M*). Error is measured as the absolute value of the truth less the estimate. (A) displays the plots by number of markers. (B) displays the exact same data but separating by method.

Unfortunately, while deconvolution performance is greatly affected by the choice of marker genes, there is generally not a known universal way to choose marker genes that will be optimal for all possible datasets. Thus, an important advantage of the hybrid approach is that its performance is consistently good for a wide range of marker genes. In Figure 6, for each dataset we plot the best-case and worst-case error of each method across all of our possible choices of *M*. We see that for five (a plurality) of the datasets (Becht, Gong, Liu, Newman PBMC and Parsons) the hybrid approach has better performance than other methods for the worst-case choice of marker genes. Furthermore, for the remaining datasets the hybrid approach has a performance within 1% of the other methods except on the Newman FL and Shen-Orr datasets (see Supplementary Figures 12 and 13). All together then, in a worst-case analysis, the hybrid approach performs better, or within 1%, of all other methods on nine out of the eleven datasets.

**FIGURE 6.**
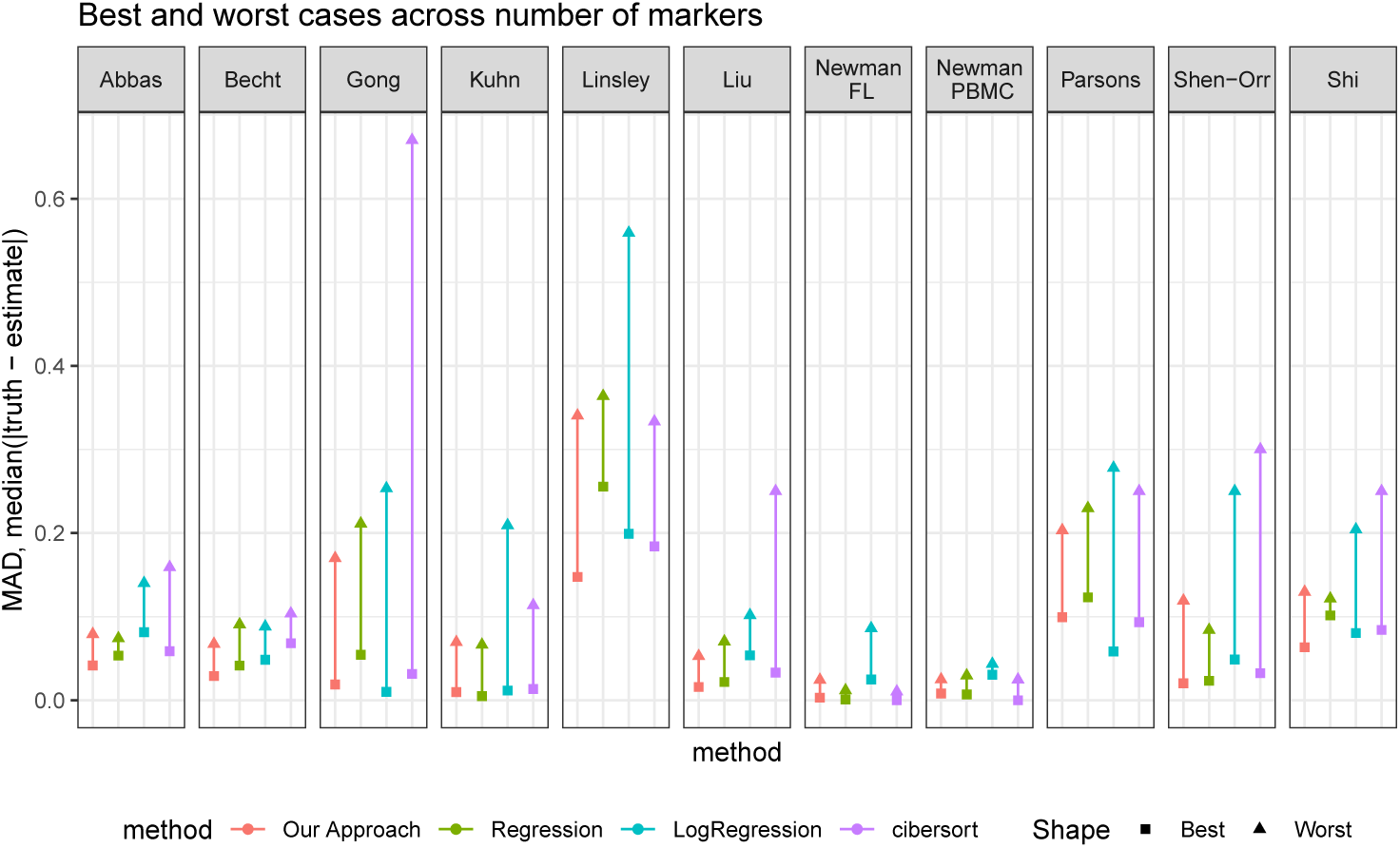
Best-case and worst-case for the methods over all datasets. The best case is the best error over all values of *M*. The worst-case is the worst error the method achieves on the dataset over all value of *M* we tried.

This worst-case analysis is important because generally there is little guidance apart from heuristics in how to choose the optimal number of marker genes. Thus, the hybrid approach has the desirable property that it will not perform too poorly if a non-optimal set of marker genes is chosen. In addition to having the best worst-case error, the hybrid approach typically has the best best-case error. In six of the eleven datasets (Abbas, Becht, Linsley, Liu, Shen-Orr, Shi) if we were to choose the optimal number of markers for each method, the hybrid approach would have the lowest error. While it is unlikely that we will choose the exact optimal markers for any dataset or method, the hybrid approach at least has the *potential* to strongly out-perform other methods. Among the remaining six datasets where the hybrid approach does not have the best best-case error, it is within 1% of the performance for all other methods except on the Parsons dataset (see Supplementary Figure 12 and 13).

## 5. CONCLUSION

Estimating cell-type proportions is an important problem with a wide range of applications across the spectrum of biological fields. Many existing deconvolution approaches estimate cell-type proportions using modified regression approaches as described in the UDAR framework. Unfortunately, fitting such a model using either linear-scale or log-scale gene expressions will be sub-optimal. Log-transforming gene expressions before fitting under a UDAR model biases the estimates. However a regression-like fit using linear-scale gene expressions assumes an un-realistic error model for gene expression measurements. Our hybrid approach tackles both of these problems proposing a model that uses a plausible mean-structure while also maintaining reasonable error assumptions. This leads to an estimate of cell-type proportions that are robust and accurate. In simulations, we saw that the hybrid approach reduces estimation variance without introducing a bias. We also saw that the model performed well under violations of Gaussian assumptions. In an analysis of real data, it was shown that cell-type deconvolution is sensitive to choice of marker genes. Unfortunately, this is compounded by the fact that for real data there is often no easy way to find an optimal set of marker genes. For this reason, the low variance estimates produced by the hybrid approach typically had the lowest error in a worst-case analysis across a range of marker genes. Furthermore, the approach also had the lowest error in a best-case analysis, showing that the variance reduction does not come at the expense of accuracy.

More broadly, the model proposed in this paper opens the door to many extensions and generalizations. While we estimate the proportions *p* in Equation 5 using a maximum-likelihood approach, one could combine this model with some of the other insights in the deconvolution literature and fit *p* using more sophisticated loss functions like L1 or L2 penalized losses, or *ε*-insensitive losses. Thus, the approach we describe has the potential to be the basis for many new, hybrid-scale, approaches to deconvolution.

In conclusion, we note that understanding cell-type heterogeneity among complex biological tissues is a problem with broad and persistent biological interest. Furthermore, an increase in high-quality cell-type reference data from bulk and single-cell sequencing technologies makes cell-type deconvolution an increasingly important tool for the analysis of high-throughput data. While historically deconvolution methods have focused on genomic data, the robust nature of the method we have proposed means it will likely be highly applicable to other high-throughput data such as methylation data or ATAC-seq. We hope to explore such directions in future work.

## 6. SOFTWARE

An implementation of the hybrid approach and examples of how to use the method can be found online at `dtangle.github.io`.

## Supporting information

Supplementary proof and plots

## SUPPLEMENTARY MATERIAL

Supplementary material includes a proof of the MLE and figures for simulation and real-data analysis.

## ACKNOWLEDGMENTS

The authors declare no conflict of interests.

## FUNDING

The authors gratefully acknowledge support from the National Science Foundation (grant no. DMS-1646108).

## Notes

https://dtangle.github.io

