## Supplementary proof and plots for "The Role of Scale in the Estimation of Cell-type Proportions"

### 1. Deriving the MLE for our Model.

We have claimed that our fitting approach described in Algorithm 2 gives the MLE for our model. We will now show that.

First we will show that maximizing the log-likelihood is equivalent to minimizing $S^2(p)$ in Algorithm 2. Consider the joint log-likelihood $\ell$ of the parameters $p$, $\sigma^2$, and $\theta$. This log-likelihood is (up to an additive constant)

$$\begin{aligned}\ell(p,\sigma^2,\theta) &= -\frac{N}{2}\log\left(\sigma^2\right) - \frac{1}{2\sigma^2}\sum_{n=1}^{N}\left(\log(Y_n) - \theta - \log\left(\sum_{k=1}^{K} p_k R_{nk}\right)\right)^2 \\ &= -\frac{N}{2}\log\left(\sigma^2\right) - \frac{1}{2\sigma^2}\sum_{n=1}^{N}\left(\lambda_n(p) - \theta\right)^2\end{aligned} \tag{1}$$

where $\lambda_n(p) \overset{def}{=} \log(Y_n) - \log\left(\sum_{k=1}^{K} p_k R_{nk}\right)$. The partial derivative with respect to $\theta$ is

$$\frac{\partial \ell}{\partial \theta} = \frac{1}{\sigma^2}\sum_{n=1}^{N}\left(\lambda_n(p) - \theta\right) \tag{2}$$

and setting this equal to zero we get the maximizing value of $\theta$ is

$$\theta^*(p) = \bar{\lambda}(p) \overset{def}{=} \frac{1}{N}\sum_{n=1}^{N}\lambda_n(p). \tag{3}$$

Furthermore, the partial derivative with respect to $\sigma^2$ is

$$\frac{\partial \ell}{\partial \sigma^2} = -\frac{N}{2\sigma^2} + \frac{1}{2\sigma^4}\sum_{n=1}^{N}\left(\lambda_n(p) - \theta\right)^2 \tag{4}$$

and setting this equal to zero we get the maximizing value for $\sigma^2$ to be

$$\sigma^{2*}(p,\theta) = \frac{1}{N}\sum_{n=1}^{N}\left(\lambda_n(p) - \theta\right)^2. \tag{5}$$

We can then form the profile likelihood for $p$ as

$$\ell^*(p) = \ell\left(p, \sigma^{2*}(p,\theta^*(p)), \theta^*(p)\right) = -\frac{N}{2}\log(\sigma^{2*}(p,\theta^*(p))) - \frac{N}{2}. \tag{6}$$

Thus we can find the MLE for $p$ by maximizing $\ell^*(p)$ or equivalently minimizing

$$\sigma^{2*}(p, \theta^*(p)) = \frac{1}{N} \sum_{n=1}^{N} \left(\lambda_n(p) - \bar{\lambda}(p)\right)^2 \tag{7}$$

which is precisely the sample variance of the $\lambda_n(p)$ defined as $S^2(p)$ in Algorithm 2.

Second, we will show that minimizing $S^2(p)$ over $[0,1]^K$ and then re-normalizing is equivalent to minimizing $S^2(p)$ over $\Delta_{K-1}$. Now if

$$p^* = \arg \min_{p \in [0,1]^K} S^2(p) \tag{8}$$

then, letting $T^* = \sum_{t=1}^{K} p_t^*$, Algorithm 2 defines $\hat{p} = p^*/T^*$. To see that $\hat{p}$ is the MLE, note that for a constant $c \in \mathbb{R}$,

$$\lambda_n(cp) = \log(Y_n) - \log\left(\sum_{k=1}^{K} cpR_{nk}\right) = \log(Y_n) - \log\left(\sum_{k=1}^{K} pR_{nk}\right) - \log(c) = \lambda_n(p) - \log(c).$$

and thus $S^2(cp) = S^2(p)$ as $S^2(p)$ is the variance of the $\lambda_n(p)$ and $S^2(cp)$ is the variance of the $\lambda_n(cp) = \lambda_n(p) - \log(c)$ and the two are equivalent since subtracting a constant $\log(c)$ does not change the sample variance $S^2$. Thus, $S^2(\hat{p}) = S^2\left(p^*/T^*\right) = S^2(p^*)$. Then since $S^2(\hat{p}) = S^2(p^*) \leq S^2(p)$ for all $p \in \Pi$ and $\Delta_{K-1} \subseteq \Pi$ then $S^2(\hat{p}) = S^2(p^*) \leq S^2(p)$ for all $p \in \Delta_{K-1}$ and so $\hat{p}$ minimizes $S^2$ over $\Delta_{K-1}$. Equivalently, $\hat{p}$ is the MLE of $p$ over $\Delta_{K-1}$.

| Name | Citation | Cell Types | Species |
|---|---|---|---|
| Shi | MAQC (2006) | 2, universal, brain | human |
| Gong | Gong *et al.* (2011) | 2, blood, breast | human |
| Shen-Orr | Shen-Orr *et al.* (2010) | 3, liver, brain, lung | rat |
| Abbas | Abbas *et al.* (2009) | 4, leukocytes | human |
| Becht | Becht *et al.* (2016) | 6, colorectal carcinoma, leukocytes | human |
| Kuhn | Kuhn *et al.* (2011) | 4, brain | rat |
| Newman FL | Newman *et al.* (2015) | 12, leukocytes | human |
| Newman PBMC | Newman *et al.* (2015) | 22, leukocytes | human |
| Parsons | Parsons *et al.* (2015) | 3, brain, liver, muscle | human |
| Liu | Liu *et al.* (2015) | 2, adenocarcinoma | human |
| Linsley | Linsley *et al.* (2014) | 3, lymphocytes,monocytes, neutrophils | human |

Table 1. Eleven benchmark deconvolution datasets.

### 2. Benchmark Datasets and Supplementary Figures

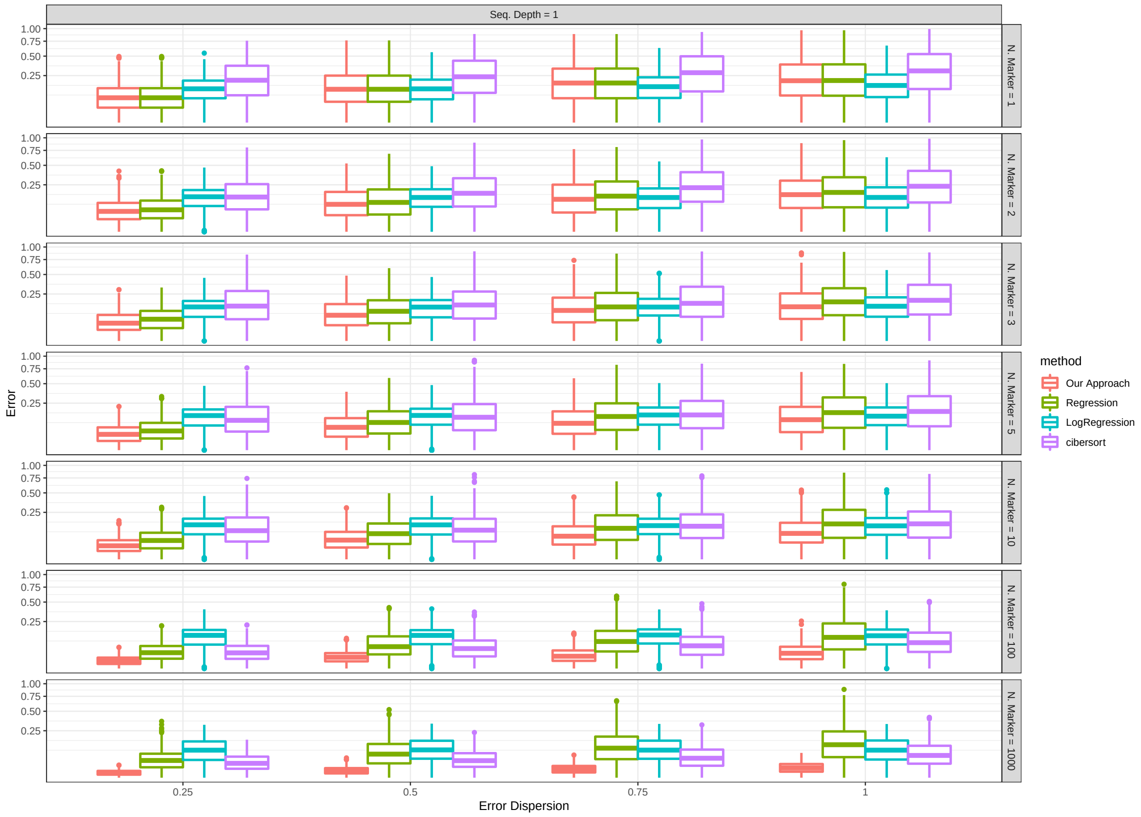


Figure 1. Evaluation of methods on simulated mixture data with Gaussian noise for different values of the variance multiplier $\tau = 1/4, 1/2, 3/4, 1$ and number of marker genes $M = 1, 2, 3, 5, 10, 100, 1000$.

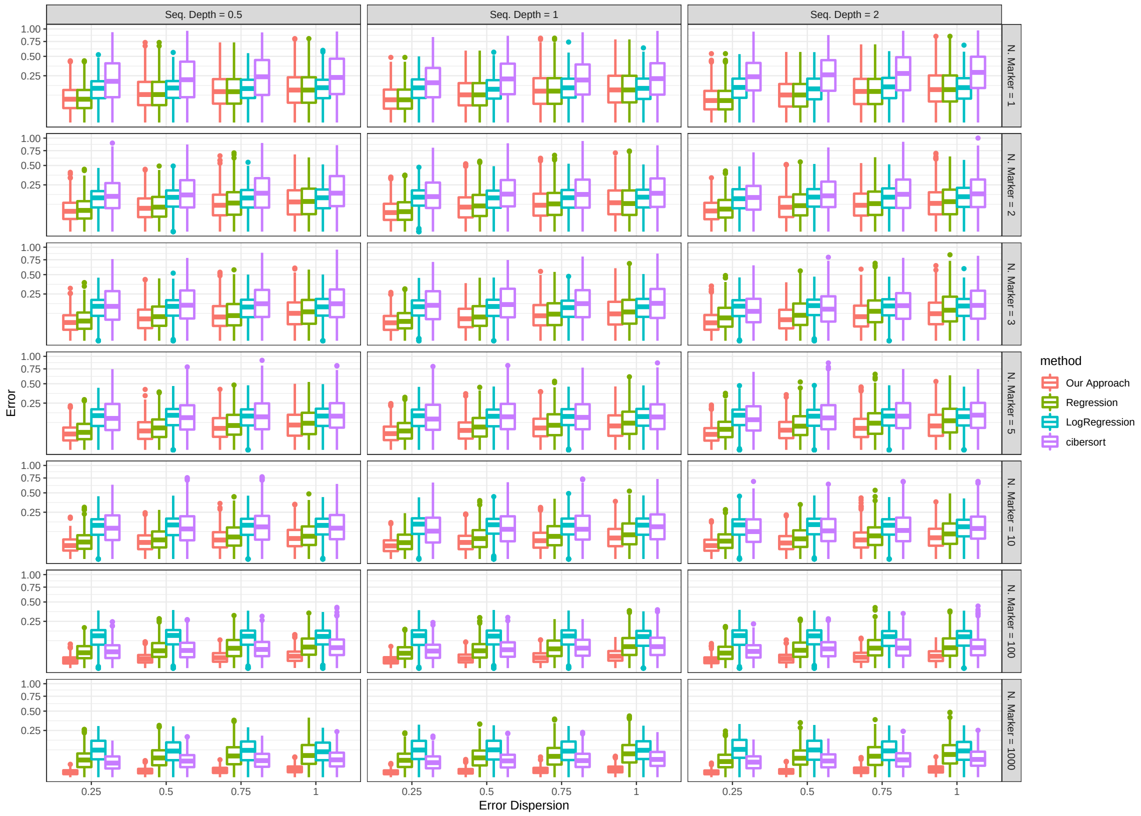


FIGURE 2. Evaluation of methods on simulated mixture data with Gaussian noise for different values of the variance multiplier $\tau = 1/4, 1/2, 3/4, 1$, number of marker genes $M = 1, 2, 3, 5, 10, 100, 1000$, and sequencing depths $1/2, 1, 2$.

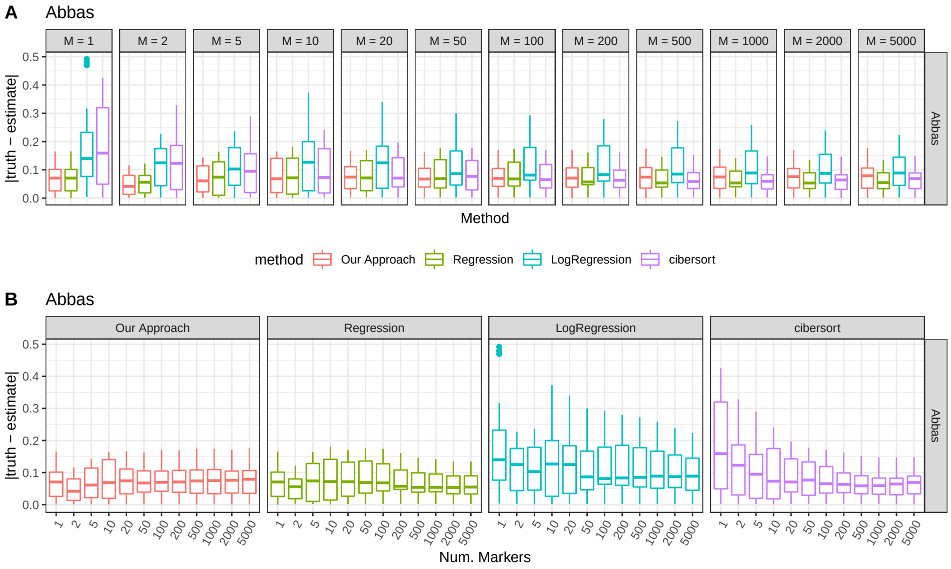


FIGURE 3. Error for methods for the Abbas dataset over a varying number of markers ($M$). Error is measured as the absolute value of the truth less the estimate. (A) displays the plots by number of markers. (B) displays the exact same data but separating by method.

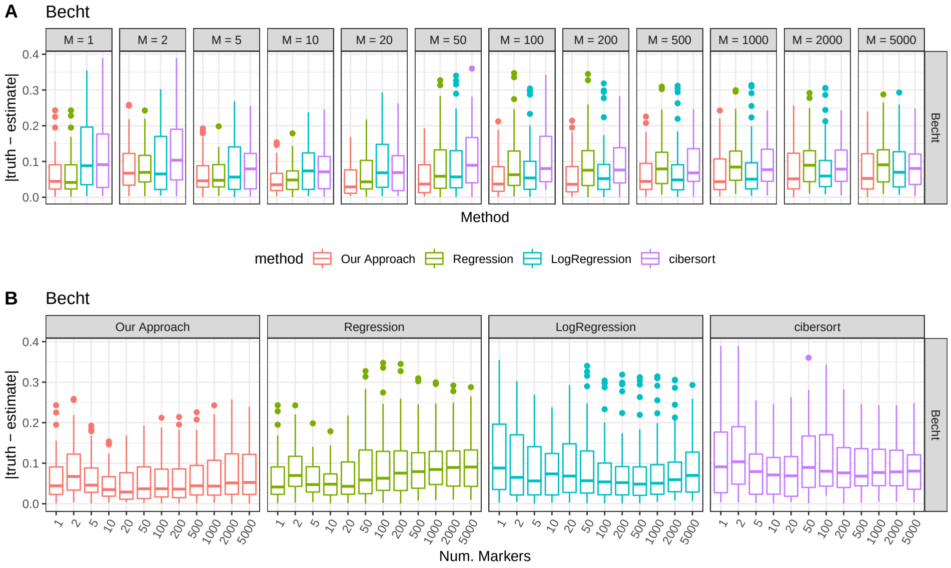


FIGURE 4. Error for methods for the Becht dataset over a varying number of markers ($M$). Error is measured as the absolute value of the truth less the estimate. (A) displays the plots by number of markers. (B) displays the exact same data but separating by method.

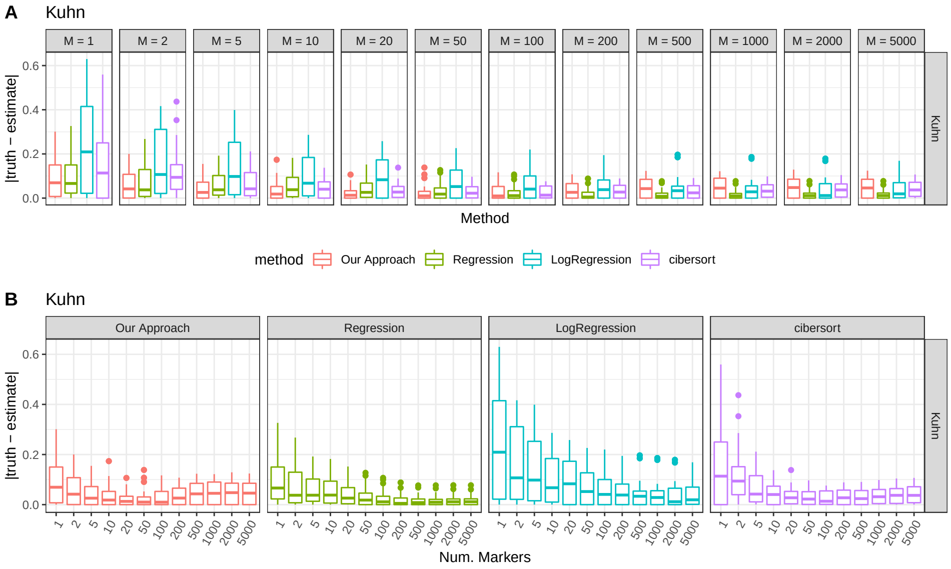


FIGURE 5. Error for methods for the Kuhn dataset over a varying number of markers ($M$). Error is measured as the absolute value of the truth less the estimate. (A) displays the plots by number of markers. (B) displays the exact same data but separating by method.

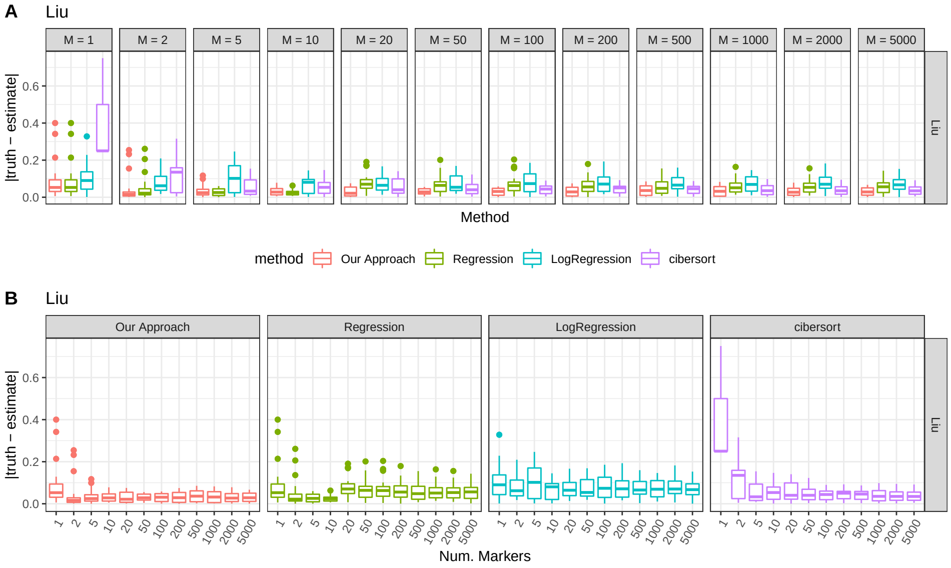


FIGURE 6. Error for methods for the Liu dataset over a varying number of markers ($M$). Error is measured as the absolute value of the truth less the estimate. (A) displays the plots by number of markers. (B) displays the exact same data but separating by method.

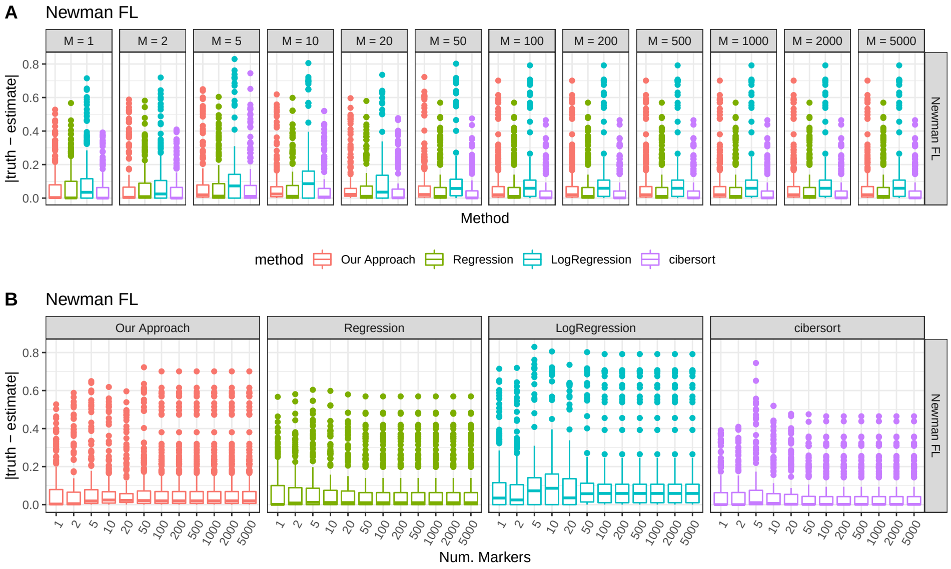


FIGURE 7. Error for methods for the Newman FL dataset over a varying number of markers ($M$). Error is measured as the absolute value of the truth less the estimate. (A) displays the plots by number of markers. (B) displays the exact same data but separating by method.

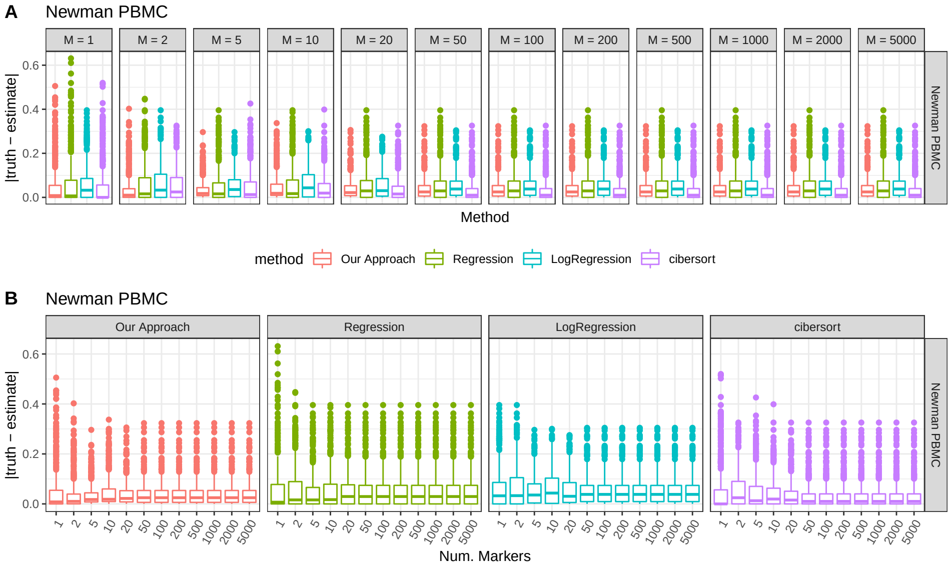


FIGURE 8. Error for methods for the Newman PBMC dataset over a varying number of markers ($M$). Error is measured as the absolute value of the truth less the estimate. (A) displays the plots by number of markers. (B) displays the exact same data but separating by method.

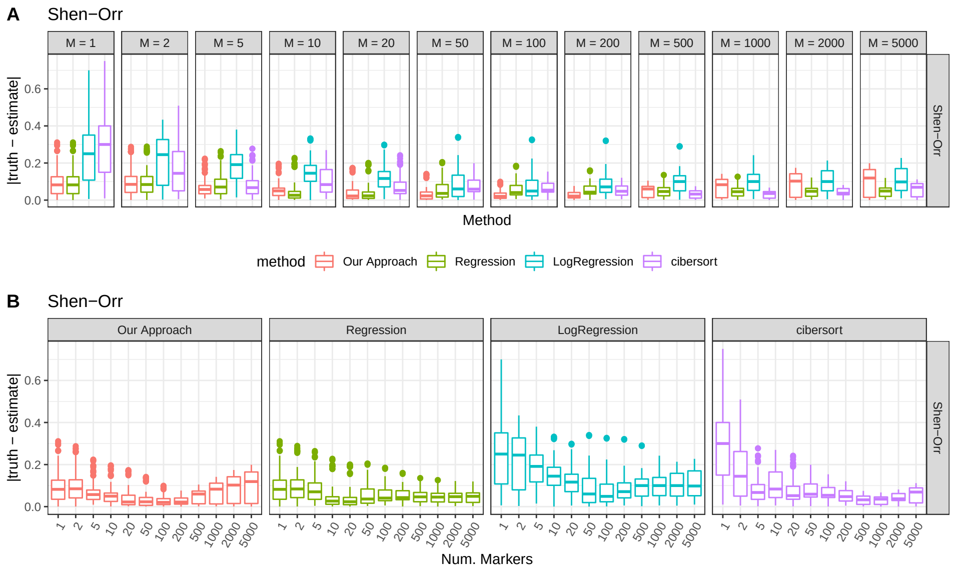


FIGURE 9. Error for methods for the Shen-Orr dataset over a varying number of markers ($M$). Error is measured as the absolute value of the truth less the estimate. (A) displays the plots by number of markers. (B) displays the exact same data but separating by method.

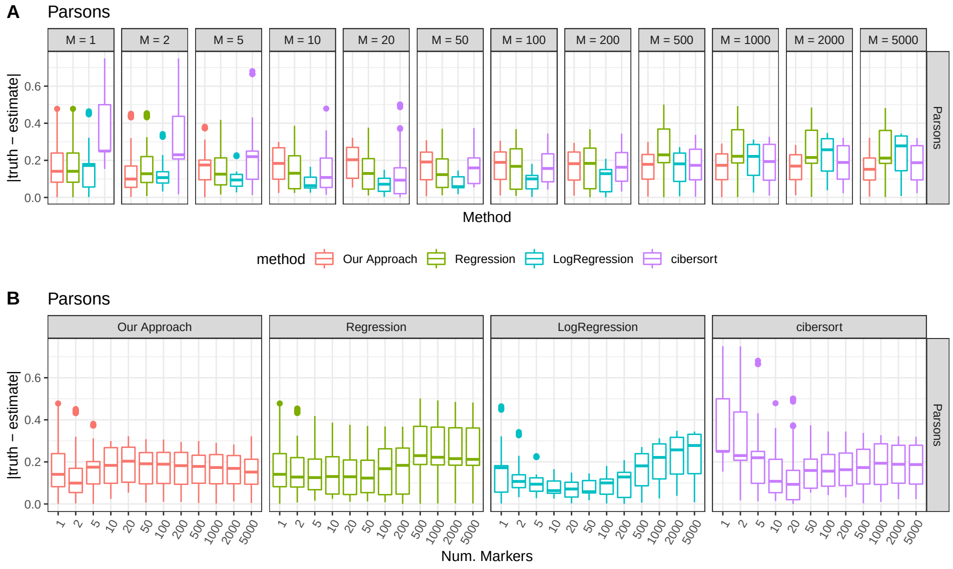


FIGURE 10. Error for methods for the Parsons dataset over a varying number of markers ($M$). Error is measured as the absolute value of the truth less the estimate. (A) displays the plots by number of markers. (B) displays the exact same data but separating by method.

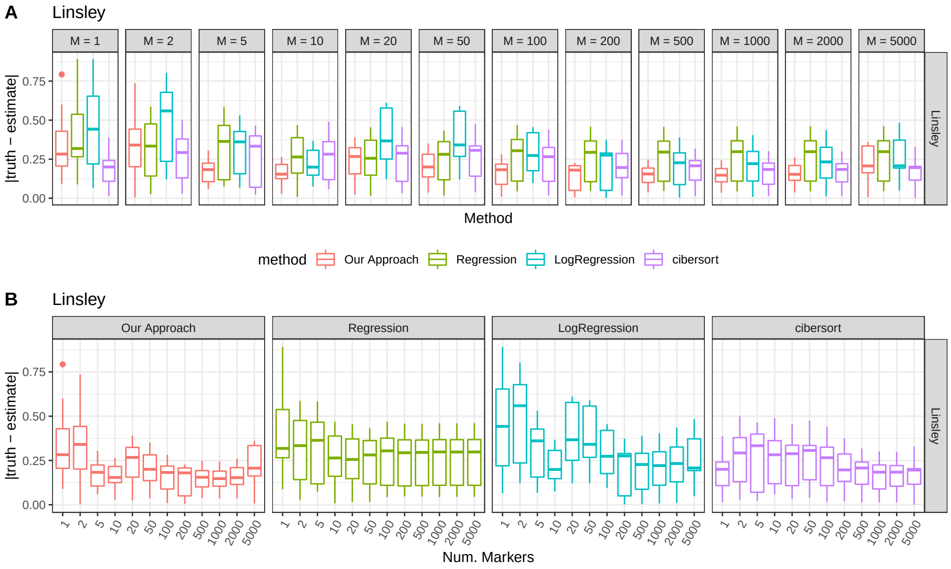


Figure 11. Error for methods for the Linsley dataset over a varying number of markers ($M$). Error is measured as the absolute value of the truth less the estimate. (A) displays the plots by number of markers. (B) displays the exact same data but separating by method.

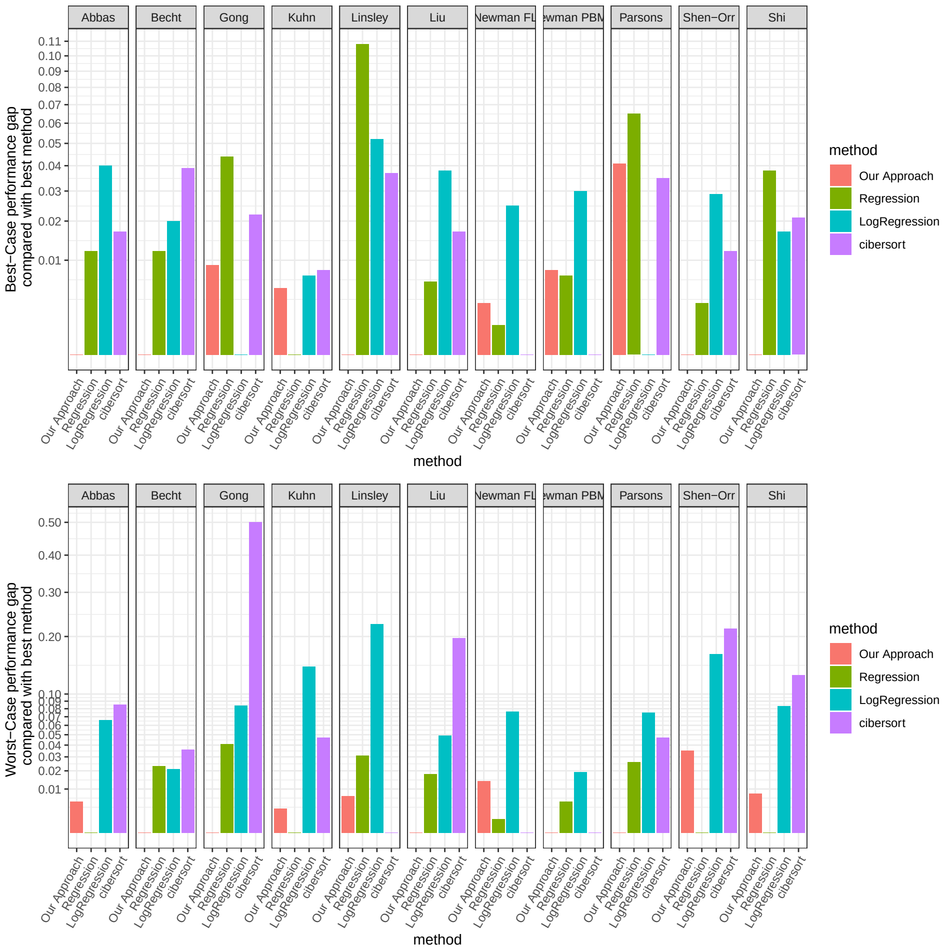


FIGURE 12. (Top) For each dataset, the y-axis displays the performance gap between the method with the lowest error and each other method. The performance gap is the difference in median accuracy for the dataset between the two methods. The performance for each method is the best-case performance for the method across the sequence of markers tested. (Bottom) Similar to the top figure but the performance for each method is the worst-case performance for the method across the sequence of markers tested.

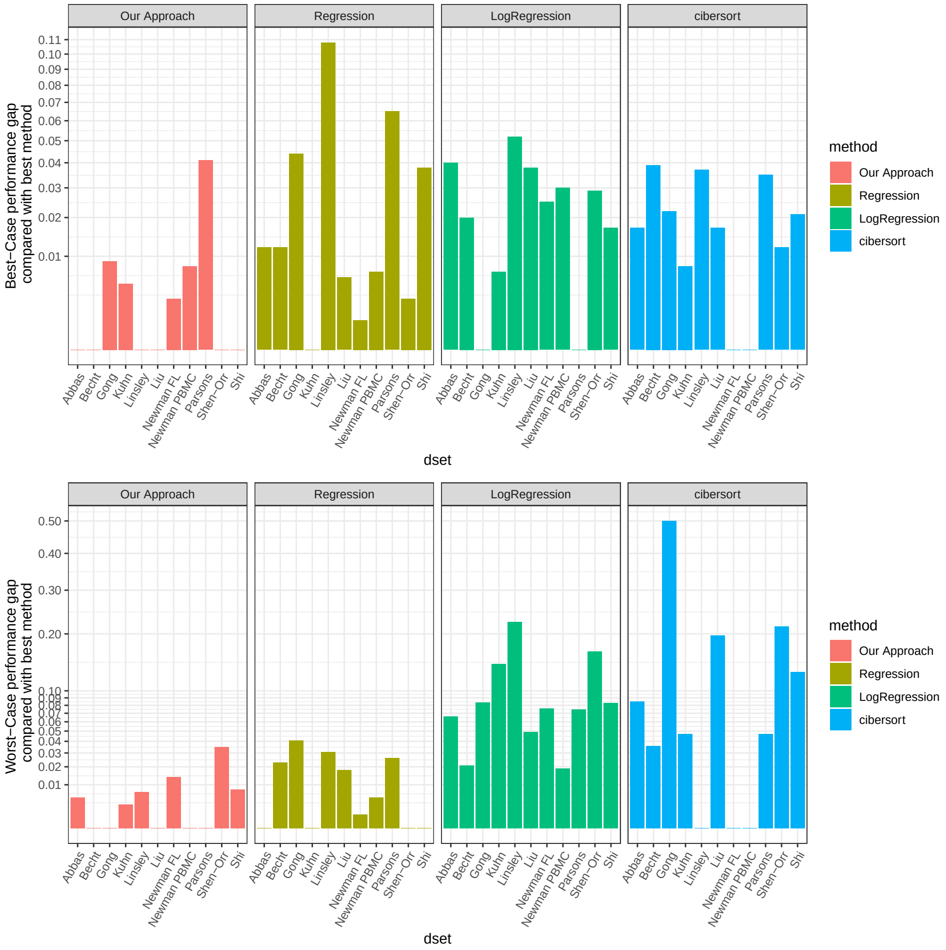


FIGURE 13. (Top) For each dataset, the y-axis displays the performance gap between the method with the lowest error and each other method. The performance gap is the difference in median accuracy for the dataset between the two methods. The performance for each method is the best-case performance for the method across the sequence of markers tested. (Bottom) Similar to the top figure but the performance for each method is the worst-case performance for the method across the sequence of markers tested.
